# Engineering acetyl-CoA metabolic shortcut for eco-friendly production of polyketides triacetic acid lactone in *Yarrowia lipolytica*

**DOI:** 10.1101/614131

**Authors:** Huan Liu, Monireh Marsafari, Fang Wang, Li Deng, Peng Xu

## Abstract

Acetyl-CoA is the central metabolic node connecting glycolysis, Krebs cycle and fatty acids synthase. Plant-derived polyketides, are assembled from acetyl-CoA and malonyl-CoA, represent a large family of biological compounds with diversified bioactivity. Harnessing microbial bioconversion is considered as a feasible approach to large-scale production of polyketides from renewable feedstocks. Most of the current polyketide production platform relied on the lengthy glycolytic steps to provide acetyl-CoA, which inherently suffers from complex regulation with metabolically-costly cofactor/ATP requirements. Using the simplest polyketide triacetic acid lactone (TAL) as a target molecule, we demonstrate that acetate uptake pathway in oleaginous yeast (*Yarrowia lipolytica*) could function as an acetyl-CoA shortcut to achieve metabolic optimality in producing polyketides. We identified the metabolic bottlenecks to rewire acetate utilization for efficient TAL production in *Y. lipolytica*, including generation of the driving force for acetyl-CoA, malonyl-CoA and NADPH. The engineered strain, with the overexpression of endogenous acetyl-CoA carboxylase (ACC1), malic enzyme (MAE1) and a bacteria-derived cytosolic pyruvate dehydrogenase (PDH), affords robust TAL production with titer up to 4.76 g/L from industrial glacier acetic acid in shake flasks, representing 8.5-times improvement over the parental strain. The acetate-to-TAL conversion ratio (0.149 g/g) reaches 31.9% of the theoretical maximum yield. The carbon flux through this acetyl-CoA metabolic shortcut exceeds the carbon flux afforded by the native acetyl-CoA pathways. Potentially, acetic acid could be manufactured in large-quantity at low-cost from Syngas fermentation or heterogenous catalysis (methanol carbonylation). This alternative carbon sources present a metabolic advantage over glucose to unleash intrinsic pathway limitations and achieve high carbon conversion efficiency and cost-efficiency. This work also highlights that low-cost acetic acid could be sustainably upgraded to high-value polyketides by oleaginous yeast species in an eco-friendly and cost-efficient manner.

## 1. Introduction

To reduce our dependence on fossil fuels and mitigate climate change concerns, researchers have started seeking alternative routes to produce fuels and commodity chemicals. Motivated by the abundant supply of renewable feedstock and agricultural waste byproducts, biochemical engineers have been able to harness various cellular metabolism to build efficient microbe cell factories with the hope to upgrade the petroleum-based chemical manufacturing process. Our increasing knowledge on microbial physiology, biochemistry and cellular genetics and sophisticated synthetic biology tools have enabled us to explore a variety of host strains with the promise to produce large volume of green and commodity chemicals, including butanol, isobutanol, 1,3-PDO, 1,4-butanediol, organic acids, amino acids, and fatty acids-based fuels *et al* in the past two decades (Zambanini, Kleineberg et al. 2016, Liu, Hu et al. 2017, Liu, Zhao et al. 2017, Awasthi, Wang et al. 2018).

Triacetic acid lactone (TAL) is an important platform chemical with a broad spectrum of industrial applications ranging from organic synthesis, polymer plasticizers, adhesives to emulsifiers. It is the precursor for the synthesis of fungicides and fine chemicals including 1,3,5-trihydroxybenzene, resorcinol, phloroglucinol *et al* (Hansen 2002). TAL is currently manufactured from chemical catalysis with petroleum-derived dehydroacetic acid followed by acidolysis with concentrated sulfuric acid (H_2_SO_4_) at high temperature (Weissermel 2008). The use of petroleum feedstock, environmentally-unfriendly catalysts and the generation of toxic byproducts limit its industrial application as green chemicals (Chia, Schwartz et al. 2012). Therefore, there is a pressing need to develop low-cost and eco-friendly platform for sustainable production of TAL from renewable feedstocks.

By manipulating acetyl-CoA and malonyl-CoA metabolism in bacteria and yeast, metabolic engineers have established a number of TAL-producing host strains with *E. coli*, *S. cerevisiae*, *Y. lipolytica et al* (Tang, Qian et al. 2013, Cardenas and Da Silva 2014, Li, Qian et al. 2018, Markham, Palmer et al. 2018). Due to cellular toxicity issues, *E. coli*, was found to be a less promising producer to accumulate TAL (Xie, Shao et al. 2006). On the contrary, *S. cerevisiae* and *Y. lipolytica* were insensitive to TAL even at concentrations up to 200 mM (25.7 g/L), opening the opportunity for us to biologically synthesize TAL in large quantity. For example, a recent study has surveyed a large panel of metabolic pathways involving energy storage and generation, pentose biosynthesis, gluconeogenesis and lipid biosynthesis, to improve TAL production in *S. cerevisiae* (Cardenas and Da Silva 2014). In another study, recycling cytosolic NADPH and limiting intracellular acetyl-CoA transportation were also found as efficient strategies to enhance TAL accumulation in S. cerevisiae (Cardenas and Da Silva 2016). Compared with *S. cerevisiae*, *Y. lipolytica*, a natural lipid producer, has recently been identified as an attractive industrial workhorse to produce various compounds due to its broad substrate range, high acetyl-CoA and malonyl-CoA flux as well as the availability of facile genetic manipulation tools (Markham and Alper 2018, Lv, Edwards et al. 2019). These features indicate that *Y. lipolytica* could be engineered as a superior host to produce acetyl-CoA and malonyl-CoA-derived compounds beyond lipid-based chemicals. Indeed, genetically rewired lipogenic pathways have achieved great success to produce lycopene, carotenoids, flavonoids and polyketides in *Y. lipolytica* (Xu, Li et al. 2014, Xu, Qiao et al. 2016, Qiao, Wasylenko et al. 2017, Xu, Qiao et al. 2017, Yu, Landberg et al. 2018). In a recent study, heterologous expression of 2-pyrone synthase along with the cytosolic expression of alternative acetyl-CoA pathways have been implemented to collectively improve TAL production up to 35.9 g/L in a bench-top reactors, with yield up to 43% of the theoretical yield from glucose (Markham, Palmer et al. 2018), highlighting the potential of using this yeast for large-scale production of polyketides.

Acetate presents in high levels in waste streams (municipal organic waste via anaerobic digestion) and lignocellulosic hydrolysates; acetate could also be synthesized from methanol carbonylation at very low cost (about $340 per metric ton) (Qian, Zhang et al. 2016). And methanol could be abundantly produced by methane oxidation with heterogenous catalysis. These facts indicate that acetic acid will be a promising, sustainable and low-cost carbon source for industrial fermentation. To systematically understand the limiting cofactors associated with polyketide production in oleaginous yeast, we use TAL biosynthetic pathway as a testbed and investigated an arrange of biological factors including cofactor (NADPH) supply, orthogonal acetyl-CoA and malonyl-CoA availability, lipogenesis competition, substrate suitability and pH conditions (Fig. 1). Stepwise optimization of these parameters led to an 8.5-times improvement of TAL production (4.76 g/L) from industrial-grade acetic acid (glacial acid), with acetic acid-to-TAL yield at 31.9% of the theoretical yield. The reported acetyl-CoA shortcut pathway delineates a strong case that we could engineer *Y. lipolytica* as an eco-friendly platform to upgrade industrial acetic acids to commodity chemicals. This work also expands our capacity to produce green chemicals from sustainable feedstocks; in particular, acetic acid could be efficiently incorporated as carbon source for cost-efficient production of various polyketides in *Y. lipolytica*.

**Fig. 1.**
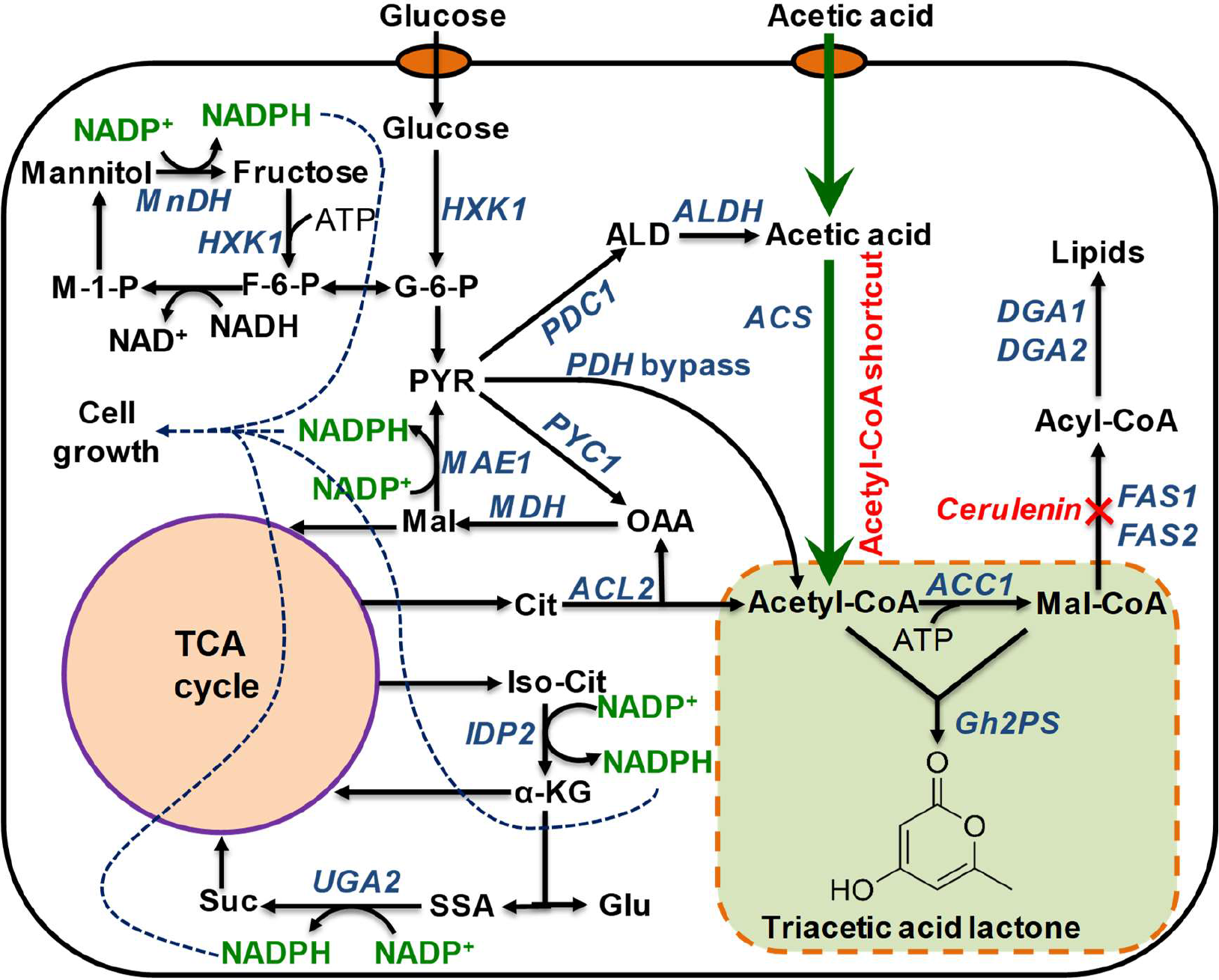
Metabolic pathway for TAL synthesis in oleaginous yeast. MnDH, mannitol dehydrogenase; HXK, hexokinase; MAE1, malic enzyme; ACL2, ATP citrate lyase; IDP2, cytosolic NADP-specific isocitrate dehydrogenase; UGA2, succinate semialdehyde dehydrogenase; PYC1, pyruvate carboxylase; PDC1, pyruvate decarboxylase; ALD, aldehyde dehydrogenase; PDH, pyruvate dehydrogenase; ACS, acetic acid synthase; FAS1 and FAS2, fatty acid synthase; ACC1, acetyl-CoA-carboxylase; GH2PS, polyketide synthase.

## 2. Materials and methods

### 2.1 Strains and cultivation

*Escherichia coli NEB5α* high efficiency competent cells were purchased from NEB for plasmid construction, preparation, propagation and storage. The *Y. lipolytica* wild type strain W29 was purchased from ATCC (ATCC20460). The auxotrophic Po1g (Leu-) was obtained from Eastern Biotech Company (Taipei, Taiwan)(Wong, Engel et al. 2017). All strains and plasmids used in this study are listed in supplementary Table S1.

LB broth or agar plate with 100 μg/mL ampicillin was used to cultivate *E. coli* strains. Yeast rich medium (YPD) was prepared with 20 g/ L Bacto peptone (Difco), 10 g/L yeast extract (Difco), and 20 g/L glucose (Sigma-Aldrich), and supplemented with 15 g/L Bacto agar (Difco) for agar plates. YNB medium with carbon/nitrogen ratio 10:1 for seed culture was made with 1.7 g/L yeast nitrogen base (without amino acids and ammonium sulfate) (Difco), 5 g/L ammonium sulfate (Sigma-Aldrich), 0.69 g/L CSM-Leu (Sunrise Science Products, Inc.) and 20 g/L glucose. Selective YNB plates were made with YNB media with carbon/nitrogen ratio 10:1 supplemented with 15 g/L Bacto agar (Difco). YNB fermentation medium with glucose as substrate and carbon/nitrogen ratio 80:1 was made with 1.7 g/L yeast nitrogen base (without amino acids and ammonium sulfate) (Difco), 1.1 g/L ammonium sulfate (Sigma-Aldrich), 0.69 g/L CSM-Leu (Sunrise Science Products, Inc.) and 40 g/L glucose.

Phosphoric buffer solution (PBS) with pH 6.0 was made with 0.2 M Na_2_HPO_4_ and 0.2 M Na_2_HPO_4_, which was used to replace water to make YNB- glucose-PBS fermentation medium. YNB fermentation medium with sodium acetate as substrate and carbon/nitrogen ratio 80:1 was made with 1.7 g/L yeast nitrogen base (without amino acids and ammonium sulfate) (Difco), 0.825 g/L ammonium sulfate (Sigma-Aldrich), 0.69 g/L CSM-Leu (Sunrise Science Products, Inc.), 41 g/L sodium acetate. Glacial acetic acids were purchased from Sigma-Aldrich. Bromocresol purple was a pH-sensitive indicator which could change its color with the pH from 5.2-7.0 (El-Ashgar, El-Basioni et al. 2012) and 40 mg/L bromocresol purple was added into fermentation medium to indicate pH variation. The pH of medium was regulated to 6.0 by 6. 0 M HCl in the fermentation process. And 1 mg/L cerulenin solution prepared with dimethylsulfoxide (DMSO) was added into fermentation medium to inhibit precursor competing pathway.

In TAL production process, single *Y. lipolytica* colonies were picked from YNB selective plates and inoculated into YNB seed liquid media, which were grown at 30 °C for 48 h. For tube test, 100 μL seed cultures were inoculated into 5 mL fermentation media in 50 mL tube. 600 μL seed cultures were inoculated into 30 mL fermentation media in 250 mL shake flasks with 250 rpm and 30 °C. Time series samples were taken for analyzing biomass, sugar content, and TAL titer.

### 2.2 Plasmid construction

All primers are listed in supplementary Table S2. All restriction enzymes were purchased from Fisher FastDigest enzymes. Plasmid miniprep, PCR clean-up, and gel DNA recoveries were using Zyppy and Zymoclean kits purchased from Zymo research. All the genes, except for *GH2PS*, were PCR-amplified with the primers from genomic DNA of *Y. lipolytica*, *S. cerevisiae*, *E. coli*, *B. subtilis*, *Aspergillus nidulans*, respectively (Supplymentary Table S1 and Table S2). GH2PS were codon-optimized according to the human codon usage table. All these genes were inserted into downstream of the *Y. lipolytica* TEF-intron promoter in the pYLXP’ vector backbone (Wong, Engel et al. 2017) at the SnaBI and KpnI site via Gibson assembly (Gibson, Young et al. 2009). Upon sequence verification by Genewiz, the restriction enzyme *Avr*II, *Nhe*I, *Not*I, *Cla*I and *Sal*I (Fermentas, Thermo Fisher Scientific) were used to digest these vectors, and the donor DNA fragments were gel purified and assembled into the recipient vector containing previous pathway components via YaliBricks subcloning protocol (Wong, Holdridge et al. 2019). All assembled plasmids were verified by gel digestion and were subsequently transformed into the *Y. lipolytica* host strain Po1g ΔLeu using the lithium acetate/single-strand carrier DNA/PEG method (Chen, Beckerich et al. 1997). In vector integration process, pYLXP’ vector assembled with functional genes was linearized by restriction enzyme NotI (Fermentas, Thermo Fisher Scientific). The linear fragment was transformed into the *Y. lipolytica* host strain Po1g ΔLeu for chromosomal gene integration.

### 2.3 Analytical methods

The cell growth was monitored by measuring the optical density at 600 nm (OD600) with a UV-vis spectrophotometer that could also be converted to dry cell weight (DCW) according to the calibration curve DCW: OD600 = 0.33:1 (g/L). The fermentation broth was centrifuged at 14,000 rpm for 5 min and the supernatant was used for analyzing the concentration of TAL, glucose, mannitol, acetic acid by HPLC with a Supelcogel ™ Carbohydrate column (Sigma). Retention times for each tested chemical were listed in Supplementary Table S3. The quantification of TAL was analyzed by an UV absorbance detector at 280 nm, while glucose, mannitol, acetic acid was examined by a refractive index detector. The column was eluted with 10 mM H_2_SO_4_ at a column temperature of 50℃, a refractive index detector at temperature of 50℃, and a flow rate of 0.4 mL/min. All chemical standards were purchased from Sigma-Aldrich and standard curve were built to derive the area-concentration relationship. All reported results are three measures with standard deviatoions.

## 3. Results and discussion

### 3.1 Boosting precursor malonyl-CoA to improve TAL production in Y. lipolytica

It has been well characterized that the plant-derived 2-pyrose synthase (2-PS) is a type III polyketide synthase (Jez, Austin et al. 2000). Specifically, 2-PS from daisy family plant *Gerbera hybrida* was found to efficiently synthesize triacetic acid lactone (TAL) by joining one acetyl-CoA with two extender malonyl-CoA molecules (Jez, Ferrer et al. 2001, Austin and Noel 2003, Xie, Shao et al. 2006). To test TAL production, the coding sequence of *G. hybrida* 2-pyrone synthase (GH2PS, Uniprot ID: P48391) was codon-optimized and overexpressed in *Y. lipolytica*, yielding 0.5 g/L of TAL at 120 h with chemically-defined complete synthetic media (CSM-leu) in test tube (Fig. 2A), demonstrating the potential of using *Y. lipolytica* as a platform to synthesize various polyketides.

**Fig. 2.**
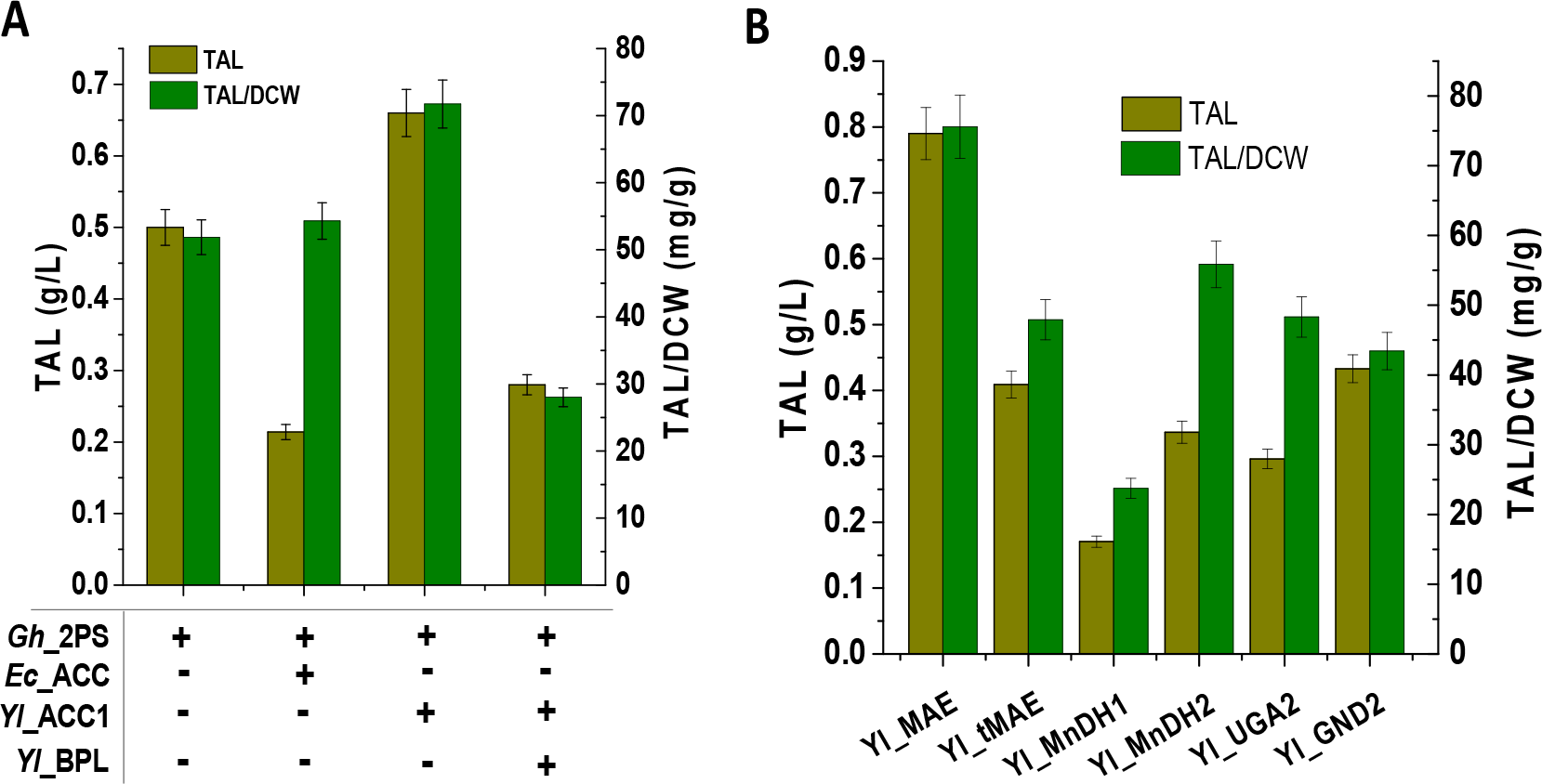
Engineering precursor pathway to improve TAL production in *Y. lipolytica*. (**A**) Comparison of TAL production from native acetyl-CoA carboxylase (ylACC1) and bacteria-derived acetyl-CoA carboxylase (EcACC). (**B**) Comparison of TAL production from alternative cytosolic NADPH pathways in *Y. lipolytica*.

It’s generally believed that malonyl-CoA is the rate-limiting precursor for polyketide synthase (Zha, Rubin-Pitel et al. 2009, Xu, Ranganathan et al. 2011). We next sought to overexpress the bacteria-derived acetyl-CoA-carboxylase and the endogenous ACC1, to test whether the conversion of acetyl-CoA to malonyl-CoA would facilitate TAL synthesis. *E. coli* intracellular malonyl-CoA could be enhanced by overexpression of the four subunits of acetyl-CoA-carboxylase (accABCD) along with the expression of the biotinylating enzyme (biotin ligase encoded by *BirA*), leading to significantly improved production of both flavonoids and fatty acids (Leonard, Lim et al. 2007, Zha, Rubin-Pitel et al. 2009, Xu, Ranganathan et al. 2011, Xu, Gu et al. 2013). With the YaliBrick gene assembly platform (Wong, Engel et al. 2017), we assembled the four catalyzing submits (accABCD) and the biotinylating subunit (*BirA*). Interestingly, a negative effect was observed when these gene clusters were introduced to *Y. lipolytica*, resulting in 40% of TAL production compared to the parental strain. These results indicate that the bacterial-derived acetyl-CoA carboxylase could not be directly translated to yeast system, presumably due to the depletion of biotins or the protein burden stress of heterologous gene expression, albeit the specific TAL production (mg/gDCW) is about 4.2% higher (Fig. 2A). Endogenous *ylACC1*, a single polypeptide encoded by YALI0C11407 with two internal introns removed, was also selected to co-express with *Gh2PS*, leading to TAL production increased to 0.66 g/L (Fig. 2A), indicating that overexpression of native ACC1 was beneficial for the TAL production. To test whether endogenous biotin ligase (BPL) was required to improve ACC1 activity (Pendini, Bailey et al. 2008), we co-expressed BPL1 (encoded by YALI0E3059). Similarly, degenerated production titer was observed in our experiment (Fig. 2A), indicating that biotin ligase is likely not the rate-limiting step for ACC1 activity in *Y. lipolytica*.

### 3.2 Recycling cytosolic NADPH to improve TAL production in Y. lipolytica

NADPH, the primary biological reducing equivalent to protect cell from oxidative stress and extend carbon-carbon backbones, has been reported as the major rate-limiting precursor in fatty acids and lipids synthesis in oleaginous species (Ratledge and Wynn 2002, Ratledge 2014, Wasylenko, Ahn et al. 2015, Qiao, Wasylenko et al. 2017). Source of cytosolic NADPH in Baker’s yeast was contributed from various alternative routes (Minard, Jennings et al. 1998, Grabowska and Chelstowska 2003, Minard and McAlister-Henn 2005), depending on the carbon source and genetic background of the yeast strain. With glucose as carbon sources, cytosolic NADPH relies on the pentose phosphate pathway. Using TAL as the target molecule, we tested a collection of auxiliary cytosolic NADPH pathways (Liu, Marsafari et al. 2019) and investigated whether these alternative pathways would improve TAL production and cellular fitness (Fig. 1). These cytosolic NADPH pathways include malic enzyme (ylMAE), mannitol dehydrogenase (ylMnDH1, ylMnDH2), 6-phosphogluconate dehydrogenase (ylGND2) and succinate semialdehyde dehydrogenase (ylUGA2), which were reported to contribute to NADPH metabolism in various fungal species (Suye and Yokoyama 1985, Minard, Jennings et al. 1998, Wynn, bin Abdul Hamid et al. 1999, Grabowska and Chelstowska 2003, Minard and McAlister-Henn 2005, Rodríguez-Frómeta, Gutiérrez et al. 2013, Liu, Marsafari et al. 2019).

Among these chosen NADPH source pathway, malic enzyme (MAE, encoded by YALI0E18634) presented the best results to improve TAL production (Fig. 2B). MAE was a NADP^+^-dependent enzyme that oxidatively decarboxylates malate to pyruvate and provides NADPH for lipid biosynthesis in oleaginous species (Ratledge 2014). When ylMAE was overexpressed with *GH2PS* and *ylACC1* (strain *HLYali101*), it showed a 60% increase in TAL titer and the volumetric production was increased to 0.8 g/L with a specific yield of 75.6 mg/g DCW (Fig. 2B). It is generally believed that ylMAE was localized in mitochondria (Zhang, Zhang et al. 2013, Ratledge 2014), we performed bioinformatic studies with TargetP 1.1 (Nielsen, Engelbrecht et al. 1997, Emanuelsson, Nielsen et al. 2000) and predicted that the first 78 nucleotides of ylMAE encodes a 26 AA leader peptide which is responsible for mitochondria targeting. We then overexpressed the truncated ylMAE (t78ylMAE) with the N-terminal MTS removed to compare how the variation of ylMAE may affect TAL synthesis. Contrary to our hypothesis, removal of the MTS signal of ylMAE exhibits adverse effect on both TAL production and cell growth (Fig. 2B). qPCR results confirmed that the ylMAE mRNA abundance was 6.8 times higher compared to the strain without ylMAE overexpression. This result indicates that the 78 nt MTS sequence is essential to the function of ylMAE. Deletion of the N-terminal 78 nt MTS sequence renders the cell with less MAE catalytic efficiency and leads to degenerated production phenotype.

Protein subcellular location in fungi has been reported to not just depend on the leader peptide sequence. For example, both *S. cerevisiae* acetyl-CoA synthase (ACS) and *Y. lipolytica* malate dehydrogenase (MDH) could move across the mitochondria or peroxisome membrane boundary to function as cytosolic enzymes, depending on the carbon source and environmental conditions (Chen, Siewers et al. 2012, Kabran, Rossignol et al. 2012, Krivoruchko, Zhang et al. 2015). In particular, one of the proven mechanisms is related with alternative splicing of mRNA transcripts in *Y. lipolytica*: where the N-terminal mRNA sequence of malate dehydrogenase could be spliced in different ways to facilitate the mature peptide targeting to distinct cellular compartment (Mekouar, Blanc-Lenfle et al. 2010, Kabran, Rossignol et al. 2012). This mechanism indicates the complicated regulation of malic enzyme in *Y. lipolytica*, highlighting that overcoming metabolite trafficking may have positive effect on TAL accumulation. Increased TAL production could also be linked to the sufficient supply of precursor malonyl-CoA, possibly due to the balanced expression of Gh2PS and ACC1, which presumably creates a metabolic sink and pushes the equilibrium of ylMAE to function in the forward direction.

### 3.3 Engineering orthogonal pyruvate dehydrogenase complex to boost TAL production in Y. lipolytica

**Fig. 3.**
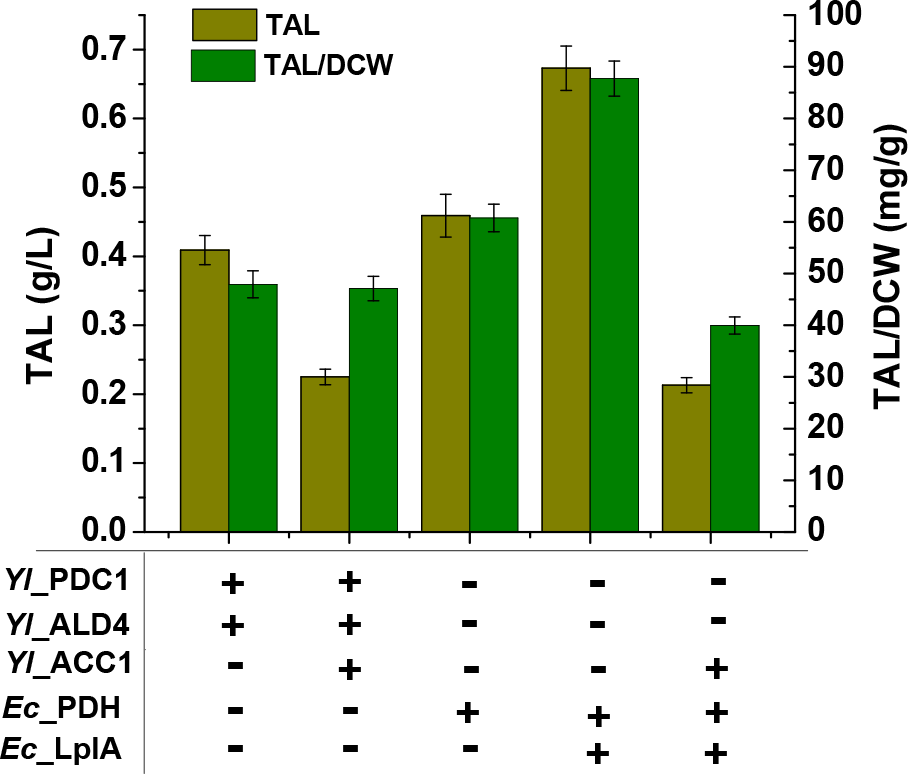
Engineering cytosolic acetyl-CoA pathway to improve TAL production in *Y. lipolytica*. YlPDC1, pyruvate decarboxylase; ylALD4, aldehyde dehydrogenase; EcPDH, pyruvate dehydrogenase comple; EcLplA, lipoate-protein ligase A.

Acetyl-CoA, derived from the central carbon metabolism (glycolysis and Krebs cycle), was the starting molecule in TAL synthesis. Acetyl-CoA is also the central hub for lipid synthesis, peroxisomal lipid oxidation and amino acid degradation pathway (Chen, Siewers et al. 2012, Krivoruchko, Zhang et al. 2015). Engineering alternative pathways to enhance cytosolic acetyl-CoA have been proven as efficient strategy to improve fatty acids and oleochemical production in both Bakers’ yeast and *Y. lipolytica* (Kozak, van Rossum et al. 2014, van Rossum, Kozak et al. 2016, Xu, Qiao et al. 2016). Here we explored two independent metabolic engineering strategies to improve the oxidation of pyruvate to cytosolic acetyl-CoA. First, we investigated the endogenous pyruvate decarboxylase (PDC), acetylaldehyde dehydrogenase (ALD) and acetyl-CoA synthase (ACS) route, which decarboxylates pyruvate to acetaldehyde by PDC, oxidizes acetaldehyde to acetate by ALD, ligates acetate with CoASH to form acetyl-CoA by ACS (Flores, Rodríguez et al. 2000). Previous results have demonstrated that we can increase lipid production when the PDC-ALD genes were overexpressed in *Y. lipolytica* (Xu, Qiao et al. 2016). Comparison of the results by combinatorially overexpressing PDCs and ALDs along with GH2PS and MAE, the best result was obtained from the strain *HLYali102*, yielding only 0.4 g/L of TAL (Fig. 3). We speculated that the synthesis process was possibly limited by the supply of malonyl–CoA. Endogenous *ylACC1* gene was further assembled to the previous pathway, however, no significant TAL improvement was observed in shake flasks (Fig. 3). These results indicate the complicated regulation of acetyl-CoA and malonyl-CoA metabolism. In particular, Snf1-mediated AMP kinase may phosphorylate ACC1 to repress ACC1 activity (Shi, Chen et al. 2014), the accumulated acetyl-CoA may lead to histone acetylation and global transcriptional change (Zhang, Galdieri et al. 2013). It could also point out a negative feedback loop of acetyl-CoA regulation: accumulated acetyl-CoA and NADH will undergo oxidative phosphorylation to generate ATP in mitochondria, which instead leads to weaker phosphorylating activity of SNF1 and elevated ACC1 activity that converts acetyl-CoA to malonyl-CoA (Mihaylova and Shaw 2011, Hsu, Liu et al. 2015).

We next attempted a bacterial-derived pyruvate dehydrogenase (PDH) complex pathway, which is orthogonal to *Y. lipolytica* metabolism. Pyruvate dehydrogenase complex (PDH), as a ubiquitous enzyme found in all kingdoms of life, catalyzed the oxidative decarboxylation of pyruvate to acetyl-CoA, forming the connection between glycolysis, the tricarboxylic acid cycle (TCA) and lipid biosynthesis (Kozak, van Rossum et al. 2014). As for all eukaryotic life including *Y. lipolytica*, PDH was localized in the mitochondrial matrix to provide acetyl-CoA for operating Krebs cycle. This mitochondrial acetyl-CoA could not be converted to TAL in the cytosol. To solve this challenge, we investigated whether the expression of the *E. coli* PDH would benefit TAL synthesis in *Y. lipolytica*. Specifically, we assembled the genes encoding the three catalytic subunits of PDH with our YaliBrick vectors (Wong, Engel et al. 2017): including pyruvate dehydrogenase (AceE, E1), dihydrolipoyllysine-residue acetyltransferase (AceF, E2), and lipoamide dehydrogenase (LpdA, E3). Nevertheless, when these three genes were assembled together and overexpressed along with GH2PS, only 0.46 g/L TAL was detected (Fig. 3). It has been discovered recently that the functional expression of bacterial PDH in *S. cerevisiae* requires the co-expression of lipoate-protein ligase A (Kozak, van Rossum et al. 2014, Kozak, van Rossum et al. 2014, Nielsen 2014), which plays a pivotal role in lipoylating E2 subunits of PDH. Similar to Baker’s yeast, the two annotated lipoyltransferase or lipoate ligase in Y. lipolytica (YALI0C03762 and YALI0F27357 respectively) are exclusively localized in mitochondria (Sulo and Martin 1993, Hermes and Cronan 2013, Cronan 2016). To enable cytosolic lipoylation and complement the bacterial PDH function, we next co-expressed the lipoate-protein ligase A (LplA), TAL production was increased to 0.67 g/L with specific yield up to 87.72 mg/gDCW (strain HLYali103) (Fig. 3 and Table S4), indicating the essential role of EcLplA as an indispensable component for the *E. coli* PDH functionality. This was the first report to functionally express the bacteria-derived EcPDH with EcLplA in *Y. lipolytica*. Creating this orthogonal acetyl-CoA route may bypass internal pathway regulations, providing a promising path to overcome metabolic limitations and improve metabolite production in *Y. lipolytica*.

Posttranslational modification of apoproteins with lipoyl groups was an important process for many enzymes. There were two complementary systems for protein lipoylation in *E. coli*: one was exogenous route using an ATP-dependent process by lipoate-protein ligase A (LplA); another one required lipoyl-[acyl-carrier protein]-protein-N-lipoyltransferase (LipB) to transfer endogenously synthesized lipoate to apoproteins (Miller, Busby et al. 2000). Both of these two proteins were localized in mitochondria, which presents a critical challenge to functionally express cytosolic PHDs in yeast. The current studies provide a viable option to functionally express bacteria-derived cytosolic PDH steps and leads to significantly improved TAL production. This PDH should be easily transferrable to other metabolite production systems, should there is a need to boost cytosolic acetyl-CoAs.

### 3.4 Comparison of TAL production from glucose and acetic acid

**Fig. 4.**
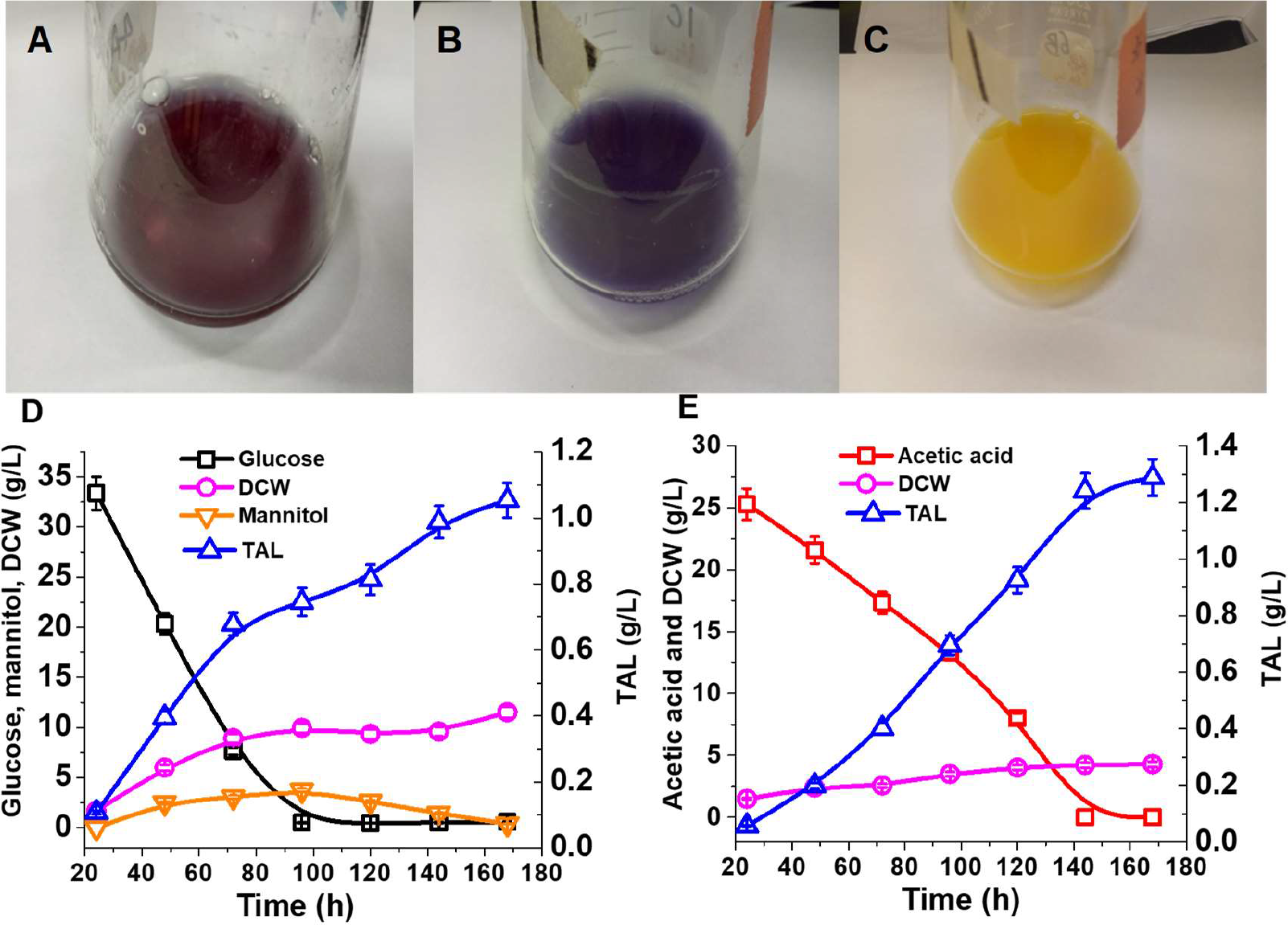
Eco-friendly production of TAL from glucose and acetic acid. (A~C) Color variation of medium with pH indicator bromocresol purple. (**A**): Initial sodium acetate (NaAc) medium with pH 6.0; (**B**): Strain cultured in NaAc medium after 48 h with pH above 8.0; (**C**): Strain cultured in glucose medium after 48 h with pH below 4.0. (**D**): TAL production from minimal media supplemented with glucose by *HLYali101*; (**E**): TAL production from minimal media supplemented with NaAc by *HLYali101*. When NaAc was used as substrate, the medium pH will increase with the consumption of acetate. We adjusted the pH to 6.0 by adding HCl. Bromocresol purple is a pH-sensitive indicator to track the pH variations of fermentation process.

A unique feature of *Y. lipolytica* is the high acetyl-CoA flux and its broad substrate range, which makes this yeast an attractive host to degrade organic waste (volatile fatty acids, alkanes *et al*). Previous metabolic engineering effort has demonstrated that *Y. lipolytica* possesses strong acetate utilization pathway that is equivalent or even superior to the hexose utilization pathway. For example, triacylglyceride (TAG) production could be improved from ~100 g/L to ~115 g/L in bench-top bioreactors, when the engineered strain was switched from glucose media (Qiao, Wasylenko et al. 2017) to acetate media (Xu, Liu et al. 2017). To investigate how different carbon sources may affect TAL synthesis, both glucose and sodium acetate were tested to compare TAL production efficiency in our engineered strains.

As acetic acid (HAc) has a relatively low pKa (4.75), we used 41 g/L sodium acetate (NaAc), equivalently to 29.5 g/L acetic acid (HAc) to cultivate the engineered strain. As the cell uptakes acetic acid, the cultivation pH will keep increasing. An *in situ* pH indicator (bromocresol purple) was used to track the pH change and 6 M HCl was used to adjust the pH in the shake flask (Fig. 4A, 4B and 4C). Controlling pH (Fig. 4A) led to the strain HLYali101 produced 1.05 g/L TAL in glucose-YNB medium, with 98% of glucose depleted and 11.53 g/L biomass produced (Fig. 4D). We also detected mannitol as a main byproduct, which increased to 3.7 g/L at 96 h and subsequently was utilized to support cell growth or TAL production (Fig. 4D). Strain HLYali101 produced about 1.29 g/L TAL from acetic acid with yield 300.69 mg/g DCW (Fig. 4E), this yield represents 230% improvement compared to the yield obtained from glucose medium (91.06 mg/g DCW) (Fig. 4D and Table S4). Depletion of acetic acid occurred at 144 h and the DCW reached 4.29 g/L, which is about half of the biomass from glucose.

Our data demonstrate that *Y. lipolytica* could efficiently uptake acetic acid as sole carbon source to produce polyketides. By harnessing the endogenous acetate uptake pathway, this externally provided acetic acid bypassed the pyruvate dehydrogenase steps and created a metabolic “shortcut” to acetyl-CoA with improved carbon conversion efficiency and pathway yield. This acetyl-CoA flux exceeded the capacity of the orthogonal PDH pathway when glucose is used as carbon source.

### 3.5 pH control and modulating precursor competing pathway to improve TAL production

**Fig. 5.**
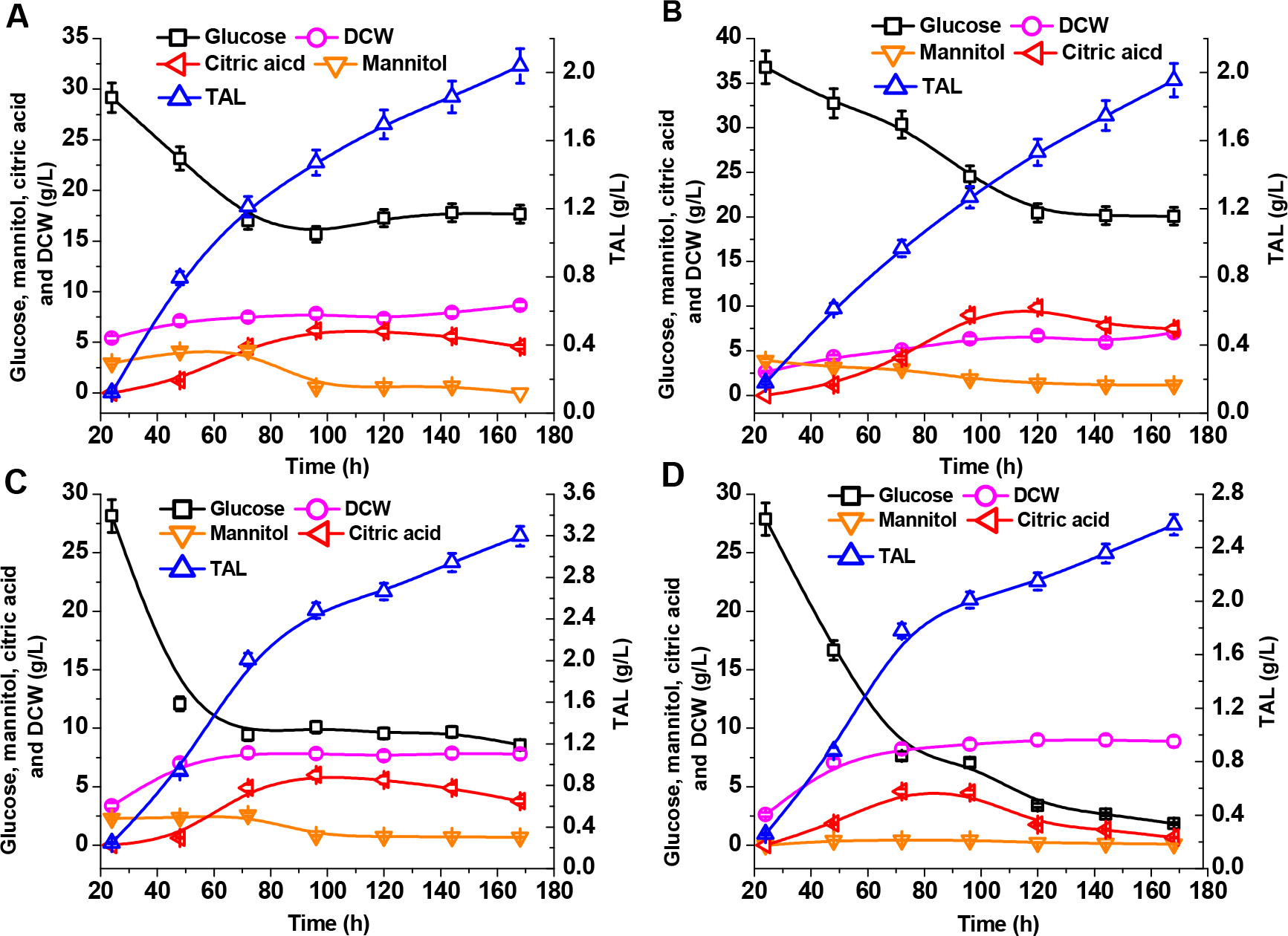
Improving TAL production by pH control and modulating precursor competing pathway in glucose-minimal media. Fermentation profile of glucose consumption, mannitol, dry cell weight, citric acid and TAL accumulation for strain *HLYali101* cultivated in glucose-minimal media conditioned with PBS buffer (**A**) or supplemented with 1 mg/L cerulenin (**C**). Fermentation profile of glucose consumption, mannitol, dry cell weight, citric acid and TAL accumulation for strain *HLYali103* cultivated in glucose-minimal media conditioned with PBS buffer (**B**) or supplemented with 1 mg/L cerulenin (**D**).

Despite that glucose is a preferred carbon source for cell growth, we constantly detected moderate or low TAL production in our engineered strains. The high TAL specific yield (300.69 mg/gDCW, Fig. 4E and Table S4) was only obtained in acetate media with pH control. We speculated that the low glucose-to-TAL yield was related with the pH declining of strain *HLYali101* (Fig. 4D): the pH of the fermentation media quickly deceased from 6 to 3.5 due to the accumulation of citric acid when glucose was utilized. This significant pH variation may negatively affect strain physiology, including cell membrane permeability and nutrients transportation process due to the loss of proton driving force. This eventually led to the accumulation of byproduct (mannitol and citric acid *et al*) and suboptimal TAL yield.

To compare how pH control may affect TAL production with the engineered strain fermenting on glucose, we supplemented the minimal YNB media with 0.2 M phosphoric buffer solution (PBS, pH 6.0). Indeed, TAL production was increased to 2.04 g/L at 168 h, which was an increase of 60.1% and 94.3% respectively, compared with the results from pH controlled (supplementary Fig. S1) and uncontrolled (Fig. 4D) process. The major byproduct mannitol reached 4.2 g/L at 72 hour and citric acid reached 6.14 g/L at 96 h, and both were subsequently utilized (Fig. 5A). The biomass of strain *HLYali101* reached 8.66 g/L DCW with the TAL specific yield at 235.6 mg/gDCW (Fig. 5A and Table S4), which is still 27.6% lower than the specific yield from acetic acid (Fig. 4E). The glucose-to-TAL yield was 91.23 mg/g, which was 3.4 times higher than that of the pH uncontrolled fermentation process (Fig. 4D and Table S4). Strain *HLYali103* (orthogonal PDH expression) led to similar fermentation profile (Fig. 5B) when the glucose YNB media was conditioned with 0.2 M PBS buffer (pH 6.0). This strain led to the accumulation of 7.01 g/L biomass with the glucose-to-TAL yield at 98.44 mg/g glucose, about 7.9% higher than the yield obtained from strain HLYali101. Similarly, only ~50% of glucose was utilized in both strain HLYali101 and HLYali103 (Fig. 5A and 5B), indicating the supplementation of PO_4_^3−^ buffer may negatively impact the glucose uptake rate.

*Y. lipolytica* is a natural lipid producer capable of accumulating 30%~60% cell weight as lipid (Xu, Qiao et al. 2017). The internal fatty acid synthase (FAS) pathway strongly competes with TAL pathway for both acetyl-CoA and malonyl-CoA. In order to mitigate FAS competition, we attempted to use FAS inhibitor (cerulenin) to suppress lipid synthesis in our engineered strain. Cerulenin could irreversibly form a covalent adduct with the active site (cysteine residue) of β-ketoacyl-ACP synthase, inhibiting the incorporation of malonyl-CoA to the extending fatty acid backbone (Vance, Goldberg et al. 1972, Price, Choi et al. 2001). When 1 mg/L cerulenin was supplemented at 48 h, 3.2 g/L TAL was produced by strain *HLYali101* at 168 h (Fig. 5C), a 57% increase compared with the result without cerulenin (Fig. 5A). Byproducts mannitol and citric acid was decreased to 0.67 g/L and 3.75 g/L at 168 h, respectively. Glucose was depleted relatively faster compared to the strain without cerulenin (Fig. 5A). And the specific TAL production reached to 363.6 mg/gDCW, a 54% increase compared to the strain without cerulenin (Fig. 5A and Table S4). Similarly, strain *HLYali103* produced about 2.57 g/L TAL (Fig. 5D), a 32% increase compared to the strain without cerulenin (Fig. 5B). These results indicated that FAS pathway was the major competing pathway and malonyl-CoA flux became the main driver of TAL synthesis in the engineered strain HLYali101 and HLYali103. Taken together, pH control and inhibition of FAS activity were effective strategies for diverting malonyl-CoA flux to TAL synthesis.

### 3.6 High-yield TAL production from acetic acid with chromosomally-integrated strains

**Fig. 6.**
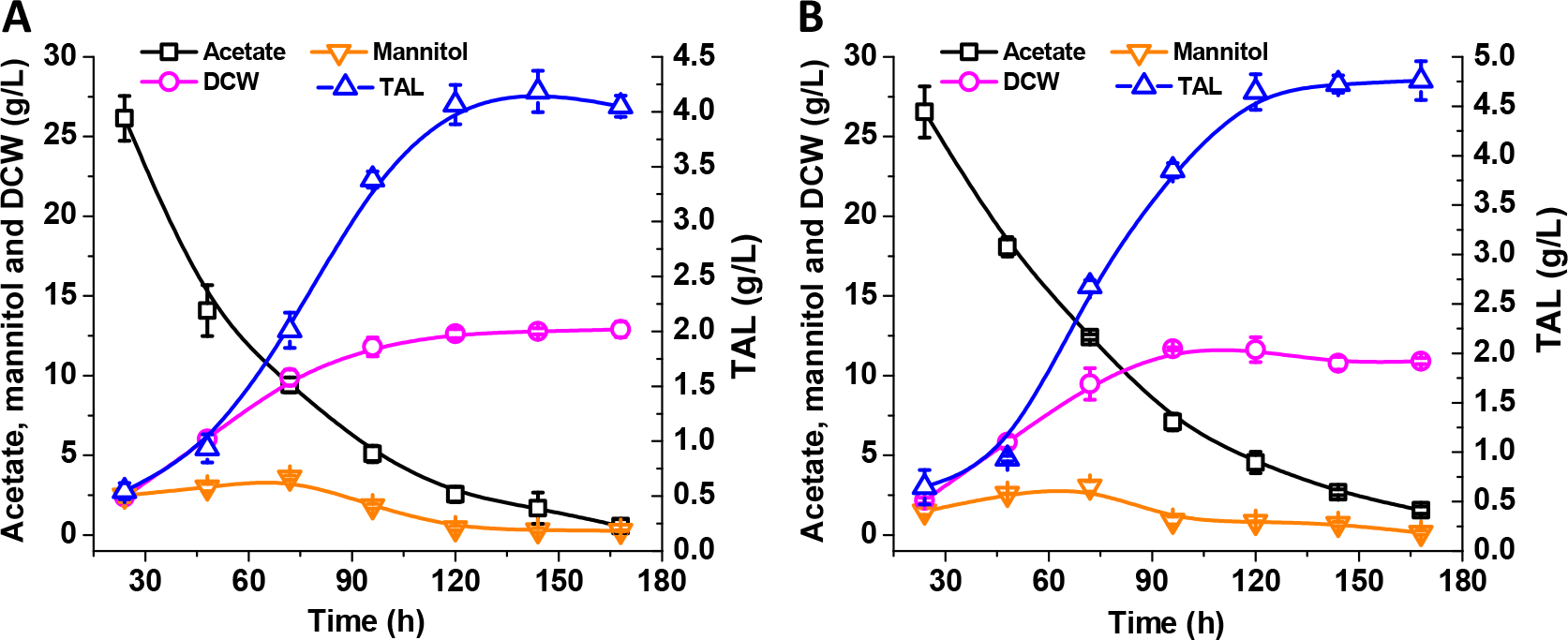
Test of TAL production with chromosomally-integrated pathway. Fermentation profile of NaAc consumption, mannitol, dry cell weight and TAL accumulation in strains LHYali105 (**A**) and LHYali106 (**B**).

The low biomass (Fig. 5C and 5D) and significant cost of cerulenin (10 mg for $272 from Sigma Aldrich) prohibits the supplementation of cerulenin as chemical inhibitors to modulate malonyl-CoA flux in the engineered strain. On the other hand, ACC1, the key metabolic node of malonyl-CoA source pathway, was primarily controlled through the phosphorylation of serine residues by Snf1-mediated AMP kinase. The overflow of glucose metabolism or nitrogen starvation activates Snf1 kinase and slows down ACC1 activity (Hedbacker and Carlson 2008, Seip, Jackson et al. 2013, Zhang, Galdieri et al. 2013, Shi, Chen et al. 2014), which makes glucose a unfavorable carbon source for the production of malonyl-CoA-derived compounds. Since acetic acid could be directly converted to acetyl-CoA, we speculate that this acetyl-CoA shortcut may present little or no activation of the Snf1 kinase. In comparison with glucose, we hypothesize this alternative carbon source (acetic acid) will not cause inhibitory effect to the malonyl-CoA source pathway (ACC1).

To increase strain stability in acetic acid, we integrated the genes Gh2PS-ylMAE1-EcPDH-EcLplA and Gh2PS-ylMAE1-ylACC1-EcPDH-EcLplA (Fig. S2) into the chromosome of *Y. lipolytica*, leading to strains LHYali105 and LHYali106, respectively. The integrated strain LHYali105 yielded about 4.18 g/L TAL from YPA complex media (yeast extract, peptone and sodium acetate) in shake flask (Fig. 6A), a 37% increase compared to the strain LHYali105 grown in glucose-based YPD (yeast extract, peptone and dextrose) media (supplementary Fig. S3). With the complementation of cytosolic PDH pathway, the strain LHYali106 yielded about 4.76 g/L TAL from YPA media (Fig. 6B), a 56% increase compared to the strain LHYali105 grown in YPD (supplementary Fig. S3). The biomass of LHYali106 reached 11.68 g/L at the end of fermentation with little mannitol (3.05 g/L) but no citric acid formation in the fermentation broth (Fig. 6B). The specific TAL yield reaches 407.5 mg/gDCW, which is 12.1% higher than the strain with cerulenin supplementation (Fig. 5C). The acetate-to-TAL conversion yield was increased to 0.149 g/g (Table S4), representing 31.9% of the theoretical maximum yield (0.467 g/g). This result demonstrates that acetic acid could lead to much higher TAL yield, exceeding the capability when glucose was used as carbon sources. Meanwhile, acetate has not been explored widely as sole carbon source in industrial fermentation. This work highlights the potential of engineering *Y. lipolytica* as a promising microbial platform for commercial manufacturing of high-value polyketides from low-cost acetic acids (glacial acetic acids).

## 4. Conclusion

In this work, triacetic acid lactone (TAL) biosynthetic pathway was constructed and optimized in oleaginous yeast *Y. lipolytica*. In addition to the codon-optimized 2-pyrone synthase gene, we found that both endogenous acetyl-CoA carboxylase (ylACC1) and malic enzyme (ylMAE1) are critical for the efficient conversion of acetyl-CoA and malonyl-CoA to TAL. Acetyl-CoA is the central hub of carbon metabolism, connecting with glycolysis, Krebs cycle and lipid synthesis. To unleash the potential of the heterologous polyketide synthase pathway, we assessed various cytosolic acetyl-CoA pathways to improve TAL synthesis. A bacteria-derived pyruvate dehydrogenase complex (Ec-PDH) along its cognate lipolyating protein (Ec-lplA) were identified as the most efficient cytosolic acetyl-CoA routes to improve TAL production. The engineered strain could efficiently ferment glucose or sodium acetate (NaAc) to TAL. In particular, the TAL titer (g/L) and specific yield (mgTAL/gDCW) from acetic acid media exceeds the titer and the specific yield obtained in glucose media. An *in situ* pH indicator (bromocresol purple) was used as a real-time pH reporter for process optimization in shake flasks. Suboptimal level of TAL accumulation (~3.2 g/L) was observed in a pH-controlled fermentation when the glucose media was supplemented with expensive FAS inhibitor (cerulenin). When the engineered strain was cultivated with acetate as sole carbon source in the absence of cerulenin, the TAL production was increased to 4.76 g/L in the chromosomally-integrated strain. The acetate-to-TAL yield reaches 0.149 g/g, representing 31.9% of the theoretical maximum yield (0.467 g/g). Since acetate uptake pathway essentially bypasses the mitochondrial PDH steps, the feeding of acetate creates a metabolic shortcut to acetyl-CoA. The carbon flux through this acetyl-CoA metabolic shortcut exceeds the carbon flux afforded by the native acetyl-CoA pathways. This alternative carbon sources present a metabolic advantage over glucose, which may not elicit or activate Snf1-mediated AMP kinase repressing ACC1 activity. Potentially, acetic acid could be a readily-available, low-cost carbon sources from Syngas fermentation, anerobic digestion or heterogenous catalysis (methanol carbonylation).

This report demonstrates that *Y. lipolytica* possesses superior acetate uptake pathway to support both cell growth and metabolite production. Our engineering strategies highlights that acetic acid could be utilized as low-cost and sustainable carbon feedstock for efficient production of plant-derived polyketides in oleaginous yeast species. More importantly, the use of metabolic shortcut may overcome many intrinsic pathway limitations (kinetic or thermodynamic limitations), and facilitate high carbon conversion efficiency and volumetric titer. Since TAL is the simplest polyketide scaffold, the reported work may also be translated to other microbial platforms for upgrading waste acetic acids to high-value compounds in an eco-friendly and cost-efficient manner.

## Acknowledgements

This work was supported by the Cellular & Biochem Engineering Program of the National Science Foundation under grant no.1805139. The authors would also like to acknowledge the Department of Chemical, Biochemical and Environmental Engineering at University of Maryland Baltimore County for funding support. HL would like to thank the China Scholarship Council for funding support.

## Author contributions

PX conceived the topic. HL and MM performed genetic engineering and fermentation experiments. HL and PX wrote the manuscript.

## References

Austin, M. B. and J. P. Noel (2003). “The chalcone synthase superfamily of type III polyketide synthases.” Natural product reports 20(1): 79–110.

Awasthi, D., L. Wang, M. S. Rhee, Q. Wang, D. Chauliac, L. O. Ingram, and K. T. Shanmugam, (2018). “Metabolic engineering of Bacillus subtilis for production of D‐lactic acid.” Biotechnol Bioeng 115(2): 453–463.

Cardenas, J. and N. A. Da Silva (2014). “Metabolic engineering of Saccharomyces cerevisiae for the production of triacetic acid lactone.” Metabolic Engineering 25: 194–203.

Cardenas, J. and N. A. Da Silva (2016). “Engineering cofactor and transport mechanisms in Saccharomyces cerevisiae for enhanced acetyl-CoA and polyketide biosynthesis.” Metabolic Engineering 36: 80–89.

Chen, D.-C., J.-M. Beckerich and C. Gaillardin (1997). “One-step transformation of the dimorphic yeast Yarrowia lipolytica.” Appl Microbiol Biotechnol 48(2): 232–235.

Chen, Y., V. Siewers and J. Nielsen (2012). “Profiling of Cytosolic and Peroxisomal Acetyl-CoA Metabolism in Saccharomyces cerevisiae.” PLOS ONE 7(8): e42475.

Chia, M., T. J. Schwartz, B. H. Shanks, and J. A. Dumesic, (2012). “Triacetic acid lactone as a potential biorenewable platform chemical.” Green Chemistry 14(7): 1850–1853.

Cronan, J. E. (2016). “Assembly of Lipoic Acid on Its Cognate Enzymes: an Extraordinary and Essential Biosynthetic Pathway.” Microbiology and Molecular Biology Reviews 80(2): 429.

El-Ashgar, N. M., A. I. El-Basioni, I. M. El-Nahhal, S. M. Zourob, T. M. El-Agez and S. A. Taya, (2012). “Sol-Gel thin films immobilized with bromocresol purple pH-sensitive indicator in presence of surfactants.” ISRN Analytical Chemistry 2012.

Emanuelsson, O., H. Nielsen, S. Brunak and G. von Heijne (2000). “Predicting Subcellular Localization of Proteins Based on their N-terminal Amino Acid Sequence.” Journal of Molecular Biology 300(4): 1005–1016.

Flores, C.-L., C. Rodríguez, T. Petit and C. Gancedo (2000). “Carbohydrate and energy-yielding metabolism in non-conventional yeasts.” FEMS microbiology reviews 24(4): 507–529.

Gibson, D. G., L. Young, R.-Y. Chuang, J. C. Venter, C. A. Hutchison, III and H. O. Smith, (2009). “Enzymatic assembly of DNA molecules up to several hundred kilobases.” Nature methods 6(5): 343.

Grabowska, D. and A. Chelstowska (2003). “The ALD6 gene product is indispensable for providing NADPH in yeast cells lacking glucose-6-phosphate dehydrogenase activity.” World J Biol Chem 278: 13984–13988.

Hansen, C. A. (2002). Chemo-enzymatic Synthesis of Aromatics Via Non-shikimate Pathway Intermediates, Michigan State University, Department of Chemistry.

Hedbacker, K. and M. Carlson (2008). “SNF1/AMPK pathways in yeast.” Front Biosci 13: 2408–2420.

Hermes, F. A. and J. E. Cronan, (2013). “The role of the Saccharomyces cerevisiae lipoate protein ligase homologue, Lip3, in lipoic acid synthesis.” Yeast 30(10): 415–427.

Hsu, H. E., T. N. Liu, C. S. Yeh, T. H. Chang, Y. C. Lo and C. F. Kao (2015). “Feedback Control of Snf1 Protein and Its Phosphorylation Is Necessary for Adaptation to Environmental Stress.” J Biol Chem 290(27): 16786–16796.

Jez, J. M., M. B. Austin, J.-L. Ferrer, M. E. Bowman, J. Schröder and J. P. Noel (2000). “Structural control of polyketide formation in plant-specific polyketide synthases.” Chemistry & Biology 7(12): 919–930.

Jez, J. M., J. L. Ferrer, M. E. Bowman, M. B. Austin, J. Schröder, R. A. Dixon, and J. P. Noel (2001). “Structure and mechanism of chalcone synthase-like polyketide synthases.” Journal of Industrial Microbiology and Biotechnology 27(6): 393–398.

Kabran, P., T. Rossignol, C. Gaillardin, J. M. Nicaud and C. Neuvéglise (2012). “Alternative splicing regulates targeting of malate dehydrogenase in Yarrowia lipolytica.” DNA Res 19(3): 231–244.

Kozak, B. U., H. M. van Rossum, K. R. Benjamin, L. Wu, J.-M. G. Daran, J. T. Pronk and A. J. A. van Maris (2014). “Replacement of the Saccharomyces cerevisiae acetyl-CoA synthetases by alternative pathways for cytosolic acetyl-CoA synthesis.” Metabolic Engineering 21: 46–59.

Kozak, B. U., H. M. van Rossum, M. A. Luttik, M. Akeroyd, K. R. Benjamin, L. Wu, S. de Vries, J.-M. Daran, J. T. Pronk and A. J. van Maris (2014). “Engineering acetyl coenzyme A supply: functional expression of a bacterial pyruvate dehydrogenase complex in the cytosol of Saccharomyces cerevisiae.” Mbio 5(5): e01696–01614.

Kozak, B. U., H. M. van Rossum, M. A. H. Luttik, M. Akeroyd, K. R. Benjamin, L. Wu, S. de Vries, J.-M. Daran, J. T. Pronk and A. J. A. van Maris (2014). “Engineering Acetyl Coenzyme A Supply: Functional Expression of a Bacterial Pyruvate Dehydrogenase Complex in the Cytosol of Saccharomyces cerevisiae Saccharomyces cerevisiae.” mBio 5(5): e01696–01614.

Krivoruchko, A., Y. Zhang, V. Siewers, Y. Chen and J. Nielsen (2015). “Microbial acetyl-CoA metabolism and metabolic engineering.” Metabolic Engineering 28: 28–42.

Krivoruchko, A., Y. Zhang, V. Siewers, Y. Chen and J. Nielsen (2015). “Microbial acetyl-CoA metabolism and metabolic engineering.” Metabolic Engineering 28(Supplement C): 28–42.

Leonard, E., K. Lim, P. Saw and M. Koffas (2007). “Engineering central metabolic pathways for high-level flavonoid production in Escherichia coli.” Applied and Environmental Microbiology: 3877–3886.

Li, Y., S. Qian, R. Dunn and P. C. Cirino (2018). “Engineering Escherichia coli to increase triacetic acid lactone (TAL) production using an optimized TAL sensor-reporter system.” J Ind Microbiol Biotechnol.

Liu, H., H. Hu, Y. Jin, X. Yue, L. Deng, F. Wang and T. Tan (2017). “Co-fermentation of a mixture of glucose and xylose to fumaric acid by Rhizopus arrhizus RH 7-13-9#.” Bioresour Technol 233: 30–33.

Liu, H., M. Marsafari, L. Deng and P. Xu (2019). “Understanding lipogenesis by dynamically profiling transcriptional activity of lipogenic promoters in Yarrowia lipolytica.” Applied Microbiology and Biotechnology.

Liu, H., S. Zhao, Y. Jin, X. Yue, L. Deng, F. Wang and T. Tan (2017). “Production of fumaric acid by immobilized Rhizopus arrhizus RH 7-13-9# on loofah fiber in a stirred-tank reactor.” Bioresour Technol 244(Part 1): 929–933.

Lv, Y., H. Edwards, J. Zhou and P. Xu (2019). “Combining 26s rDNA and the Cre-loxP system for iterative gene integration and efficient marker curation in Yarrowia lipolytica.” ACS Synthetic Biology.

Markham, K. A. and H. S. Alper (2018). “Synthetic Biology Expands the Industrial Potential of Yarrowia lipolytica.” Trends in biotechnology.

Markham, K. A., C. M. Palmer, M. Chwatko, J. M. Wagner, C. Murray, S. Vazquez, A. Swaminathan, I. Chakravarty, N. A. Lynd and H. S. Alper (2018). “Rewiring Yarrowia lipolytica toward triacetic acid lactone for materials generation.” Proceedings of the National Academy of Sciences: 201721203.

Mekouar, M., I. Blanc-Lenfle, C. Ozanne, C. Da Silva, C. Cruaud, P. Wincker, C. Gaillardin and C. Neuvéglise (2010). “Detection and analysis of alternative splicing in Yarrowia lipolytica reveal structural constraints facilitating nonsense-mediated decay of intron-retaining transcripts.” Genome Biology 11(6): R65–R65.

Mihaylova, M. M. and R. J. Shaw (2011). “The AMPK signalling pathway coordinates cell growth, autophagy and metabolism.” Nature cell biology 13(9): 1016–1023.

Miller, J. R., R. W. Busby, S. W. Jordan, J. Cheek, T. F. Henshaw, G. W. Ashley, J. B. Broderick, J. E. Cronan and M. A. Marletta (2000). “Escherichia coli LipA is a lipoyl synthase: in vitro biosynthesis of lipoylated pyruvate dehydrogenase complex from octanoyl-acyl carrier protein.” Biochemistry 39(49): 15166–15178.

Minard, K. I., G. T. Jennings, T. M. Loftus, D. Xuan and L. McAlister-Henn (1998). “Sources of NADPH and expression of mammalian NADP+-specific isocitrate dehydrogenases in Saccharomyces cerevisiae.” J Biol Chem 273(47): 31486–31493.

Minard, K. I. and L. McAlister-Henn (2005). “Sources of NADPH in yeast vary with carbon source.” J Biol Chem 280: 39890–39896.

Nielsen, H., J. Engelbrecht, S. Brunak and G. von Heijne (1997). “Identification of prokaryotic and eukaryotic signal peptides and prediction of their cleavage sites.” Protein Eng 10(1): 1–6.

Nielsen, J. (2014). “Synthetic Biology for Engineering Acetyl Coenzyme A Metabolism in Yeast.” mBio 5(6).

Pendini, N. R., L. M. Bailey, G. W. Booker, M. C. Wilce, J. C. Wallace and S. W. Polyak (2008). “Microbial biotin protein ligases aid in understanding holocarboxylase synthetase deficiency.” Biochimica et Biophysica Acta (BBA) - Proteins and Proteomics 1784(7): 973–982.

Price, A. C., K.-H. Choi, R. J. Heath, Z. Li, S. W. White and C. O. Rock (2001). “Inhibition of β-ketoacyl-acyl carrier protein synthases by thiolactomycin and cerulenin structure and mechanism.” Journal Of Biological Chemistry 276(9): 6551–6559.

Qian, Q., J. Zhang, M. Cui and B. Han (2016). “Synthesis of acetic acid via methanol hydrocarboxylation with CO2 and H2.” Nature Communications 7: 11481.

Qiao, K., T. M. Wasylenko, K. Zhou, P. Xu and G. Stephanopoulos (2017). “Lipid production in Yarrowia lipolytica is maximized by engineering cytosolic redox metabolism.” Nat Biotechnol 35(2): 173–177.

Ratledge, C. (2014). “The role of malic enzyme as the provider of NADPH in oleaginous microorganisms: a reappraisal and unsolved problems.” Biotechnology Letters 36(8): 1557–1568.

Ratledge, C. and J. P. Wynn (2002). “The biochemistry and molecular biology of lipid accumulation in oleaginous microorganisms.” Adv Appl Microbiol 51.

Rodríguez-Frómeta, R. A., A. Gutiérrez, S. Torres-Martínez and V. Garre (2013). “Malic enzyme activity is not the only bottleneck for lipid accumulation in the oleaginous fungus Mucor circinelloides.” Applied Microbiology and Biotechnology 97(7): 3063–3072.

Seip, J., R. Jackson, H. He, Q. Zhu and S.-P. Hong (2013). “Snf1 Is a Regulator of Lipid Accumulation in Yarrowia lipolytica.” Applied and Environmental Microbiology 79(23): 7360–7370.

Shi, S., Y. Chen, V. Siewers and J. Nielsen (2014). “Improving Production of Malonyl Coenzyme A-Derived Metabolites by Abolishing Snf1-Dependent Regulation of Acc1.” mBio 5(3).

Sulo, P. and N. C. Martin (1993). “Isolation and characterization of LIP5. A lipoate biosynthetic locus of Saccharomyces cerevisiae.” Journal of Biological Chemistry 268(23): 17634–17639.

Suye, S. and S. Yokoyama (1985). “NADPH production from NADP+ using malic enzyme of Achromobacter parvulus IFO-13182.” Enzyme and Microbial Technology 7(9): 418–424.

Tang, S.-Y., S. Qian, O. Akinterinwa, C. S. Frei, J. A. Gredell and P. C. Cirino (2013). “Screening for enhanced triacetic acid lactone production by recombinant Escherichia coli expressing a designed triacetic acid lactone reporter.” Journal Of the American Chemical Society 135(27): 10099–10103.

van Rossum, H. M., B. U. Kozak, J. T. Pronk and A. J. A. van Maris (2016). “Engineering cytosolic acetyl-coenzyme A supply in Saccharomyces cerevisiae: Pathway stoichiometry, free-energy conservation and redox-cofactor balancing.” Metabolic Engineering 36: 99–115.

Vance, D., I. Goldberg, O. Mitsuhashi, K. Bloch, S. Ōmura and S. Nomura (1972). “Inhibition of fatty acid synthetases by the antibiotic cerulenin.” Biochemical And Biophysical Research Communications 48(3): 649–656.

Wasylenko, T. M., W. S. Ahn and G. Stephanopoulos (2015). “The oxidative pentose phosphate pathway is the primary source of NADPH for lipid overproduction from glucose in Yarrowia lipolytica.” Metab Eng 30: 27–39.

Weissermel, K. (2008). Industrial organic chemistry, John Wiley & Sons.

Wong, L., J. Engel, E. Jin, B. Holdridge and P. Xu (2017). “YaliBricks, a versatile genetic toolkit for streamlined and rapid pathway engineering in Yarrowia lipolytica.” Metabolic engineering communications 5: 68–77.

Wong, L., B. Holdridge, J. Engel and P. Xu (2019). Genetic Tools for Streamlined and Accelerated Pathway Engineering in Yarrowia lipolytica. Microbial Metabolic Engineering: Methods and Protocols.

C. N. S. Santos and P. K. Ajikumar. New York, NY, Springer New York: 155–177.

Wynn, J. P., A. bin Abdul Hamid and C. Ratledge (1999). “The role of malic enzyme in the regulation of lipid accumulation in filamentous fungi.” Microbiology 145 (Pt 8): 1911–1917.

Xie, D., Z. Shao, J. Achkar, W. Zha, J. W. Frost and H. Zhao (2006). “Microbial synthesis of triacetic acid lactone.” Biotechnology and Bioengineering 93(4): 727–736.

Xie, D., Z. Shao, J. Achkar, W. Zha, J. W. Frost and H. Zhao (2006). “Microbial synthesis of triacetic acid lactone.” Biotechnol Bioeng 93(4): 727–736.

Xu, J., N. Liu, K. Qiao, S. Vogg and G. Stephanopoulos (2017). “Application of metabolic controls for the maximization of lipid production in semicontinuous fermentation.” Proceedings of the National Academy of Sciences 114(27): E5308–E5316.

Xu, P., Q. Gu, W. Wang, L. Wong, A. G. W. Bower, C. H. Collins and M. A. G. Koffas (2013). “Modular optimization of multi-gene pathways for fatty acids production in E. coli.” Nature Communications 4: 1409.

Xu, P., L. Li, F. Zhang, G. Stephanopoulos and M. Koffas (2014). “Improving fatty acids production by engineering dynamic pathway regulation and metabolic control.” Proceedings of the National Academy of Sciences 111(31): 11299–11304.

Xu, P., K. Qiao, W. S. Ahn and G. Stephanopoulos (2016). “Engineering Yarrowia lipolytica as a platform for synthesis of drop-in transportation fuels and oleochemicals.” Proceedings of the National Academy of Sciences 113(39): 10848–10853.

Xu, P., K. Qiao and G. Stephanopoulos (2017). “Engineering oxidative stress defense pathways to build a robust lipid production platform in Yarrowia lipolytica.” Biotechnol Bioeng 114(7): 1521–1530.

Xu, P., S. Ranganathan, Z. Fowler, C. Maranas and M. Koffas (2011). “Genome-scale metabolic network modeling results in minimal interventions that cooperatively force carbon flux towards malonyl-CoA.” Metabolic Engineering 13(5): 578–587.

Yu, J., J. Landberg, F. Shavarebi, V. Bilanchone, A. Okerlund, U. Wanninayake, L. Zhao, G. Kraus and S. Sandmeyer (2018). “Bioengineering triacetic acid lactone production in Yarrowia lipolytica for pogostone synthesis.” Biotechnol Bioeng.

Zambanini, T., W. Kleineberg, E. Sarikaya, J. M. Buescher, G. Meurer, N. Wierckx and L. M. Blank (2016). “Enhanced malic acid production from glycerol with high-cell density Ustilago trichophora TZ1 cultivations.” Biotechnology for Biofuels 9(1): 135.

Zha, W., S. B. Rubin-Pitel, Z. Shao and H. Zhao (2009). “Improving cellular malonyl-CoA level in Escherichia coli via metabolic engineering.” Metabolic Engineering 11(3): 192–198.

Zhang, H., L. Zhang, H. Chen, Y. Q. Chen, C. Ratledge, Y. Song and W. Chen (2013). “Regulatory properties of malic enzyme in the oleaginous yeast, Yarrowia lipolytica, and its non-involvement in lipid accumulation.” Biotechnology Letters 35(12): 2091–2098.

Zhang, M., L. Galdieri and A. Vancura (2013). “The Yeast AMPK Homolog SNF1 Regulates Acetyl Coenzyme A Homeostasis and Histone Acetylation.” Molecular and Cellular Biology 33(23): 4701–4717.

