## Supplemental figures and tables for "Engineering acetyl-CoA metabolic shortcut for eco-friendly production of polyketides triacetic acid lactone in *Yarrowia lipolytica*"

### 1. pH control to improve TAL production in glucose media

pH was known as an important factor for all the microorganisms in fermentation process, which not only influenced the morphogenic action, but also had a great effect on the growth of yeast(Szabo and Štofaníková 2002). It was reported that external pH and nitrogen sources were two critical aspects for cellular morphology and cell growth regulation in *Y. lipolytica*(Szabo 1999). The acid (AXP) and alkaline (AEP) extracellular proteases were two proteolytic enzymes providing transportable organic sources of carbon and nitrogen to support cell growth in*Y. Lipolytica*, and the expression of them were regulated by pH Polyketide synthase from *Rheum palmatum* as a crucial role in the biosynthesis of phenylbutanones has been studied in the effect of pH with an optimum range of 8.0-8.8(Abe, Takahashi et al. 2001). It has also been proven that pH control was beneficial for polyketide synthesis such as erythromycin(Elmahdi, Baganz et al. 2003).

To compare how pH control may affect TAL production with the engineered strain fermenting on glucose, we thus conducted a same strategy that the pH in the fermentation process of glucose was regulate every 24 h in shake flasks and the TAL production result was shown in Supplementary Fig. S1. The highest production of TAL reached 1.28 g/L at 144 h, which was increased by 20% compared with the pH uncontrolled fermentation process. Furthermore, glucose consumption rate became faster and it was almost used up at 72 h, but the DCW finally became lower. For byproduct, we observed that the highest titer of mannitol was half of it in pH uncontrolled process, while a new byproduct citric acid was detected in the broth which was up up to 6.5 g/L at 96 h and reduced to 3.15 g/L at the end of fermentation. It indicated that a low pH led a favorable condition for the synthesis of mannitol and inhibited the formation of citric acid due to acidic environment. From these results, we made sure that pH was an essential fermentation factor for TAL synthesis.

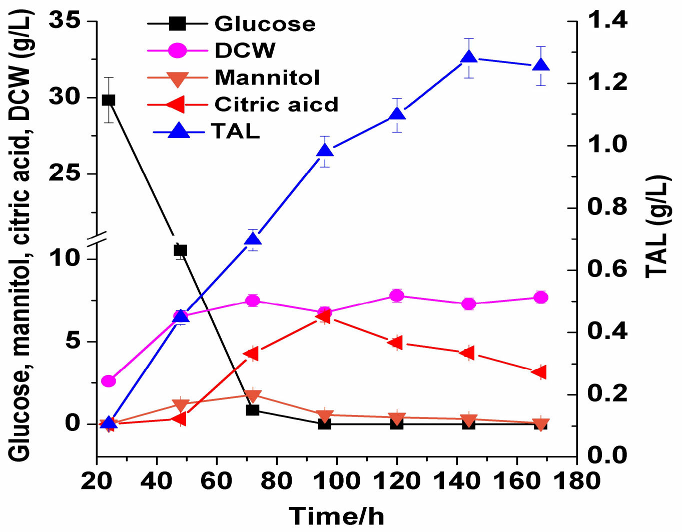


Fig. S1 TAL production by *HLYali101* with pH regulated by HCL.

### 2. Plasmid construction of PYLXP'-GH_2PS- yl MAE-yl ACC1-Ec PDH

To enhance TAL synthesis, we assembled ylACC1 gene with the previous PDH vectors (supplementary Fig. S1). When this 30.4 kb pathway were introduced to P01g and all the genes were expressed on an episomal plasmid, the growth of the transformed strain (HLYali104) was negatively impacted, and we didn't get further improvement in the TAL production, possibly due to the protein burden effect associated with the expression of large gene clusters in this yeast, but it could function when it was chromosomally-integrated.

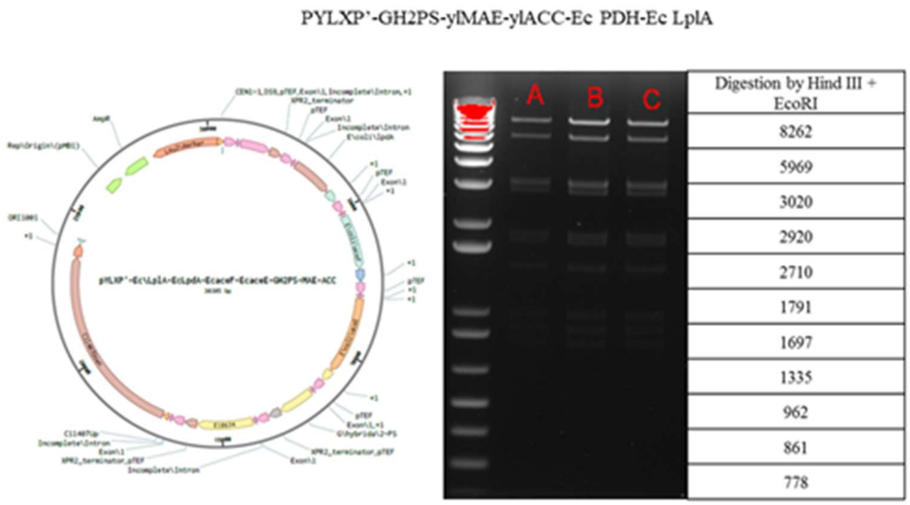


Fig. S2 Construction of vector by assembling GH2PS, ylMAE, ylACC1 with EcPDH, LplA.

### 3. TAL production by *HLYali105* from YPD medium

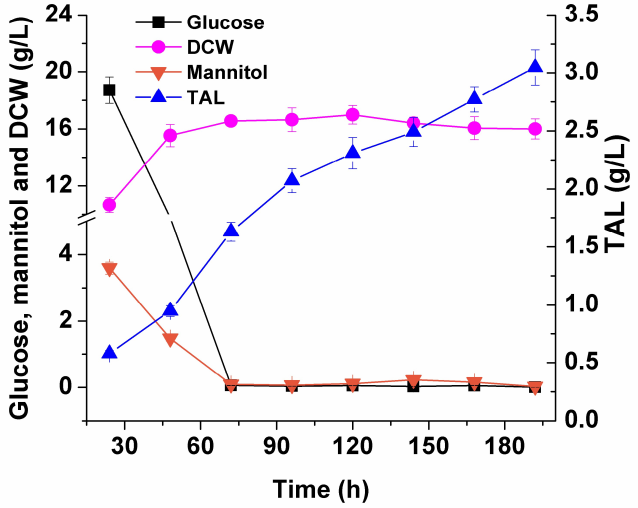


Fig. S3 TAL production from YPD medium by strain *HLYali105*

0.01M Phosphoric buffer solution (PBS) with pH 6.0was used to prepare YPD medium for Fed-batch fermentation. To maintain the dissolved oxygen levels above 20%, the agitation speed was controlled at 400-800 RPM. Sparged air was supplied at 1vvm. The initial pH of medium was 6.0 and it was base controlled using a 4 M NaOH solution and 2 M HCL solution. Three separate samples were taken every 9 or 15 hours, which would be used to detect dry cell weight (DCW), glucose consumption, TAL and by-product titer.

The vector PYLXP' containing genes *GH2PS, ylMAE*, *Ec PDH*, *Ec LplA* was integrated into genome DNA of *Y.lipolytica* to investigate the TAL production from YPD medium by the genetic-stable strain. After cultured in YPD medium for two times and selected in glucose YNB medium, the genetic-stable strain *HLYali105* was selected from eight colonies, which could produce 1 g/L TAL in tube culture. Then strain *HLYali105* was investigated on TAL accumulation in YPD medium and the results were shown in Fig. S3. It showed that the highest TAL production reached 3.05 g/L. It indicated that the genetic-stable strain *HLYali105* cultured in YDP was significant in TAL accumulation process in YNB medium. It was interesting that there was no by-product citric acid detected in this process, while another one mannitol was up to 3.59 g/L at 24 h and then converted by the strain. It was supposed that there were a lot of nutriments in YPD medium especially NADPH/NADH, which was enough for strain to grow so that glucose could be efficiently converted to TAL and biomass. Therefore, glucose was almost used up and biomass of the strain was so high, about 16 g/L. It illustrated that amount of carbon source flow into Kreb's cycle which meant that the conversion efficiency of carbon source to TAL was low. It was supposed that a high viscosity and density of broth, a low dissolved oxygen and unregulated pH could influence TAL accumulation process. Fermentation process was thus conducted in a 3 L stirred tank bioreactor to enhance TAL accumulation, but a low titer of TAL 3.89 g/L was gotten at 178 h, which meant that the strain has a potential utilization for TAL production while a further conditions optimization should be explored.

**Table S1. Strains and plasmids used in this study.**

| Plasmid or strain | Relevant properties or genotype | Source |
|---|---|---|
| **Plasmids** | | |
| PYLXP' | pYaliA1 vector backbone with leucine marker and Ampicillin resistance gene | (Xu, Qiao et al. 2017) |
| PYLXP'-GH2PS | PYLXP' carrying GH2PS from *Gerbera hybrida* | This study |
| PYLXP'-GH2PS-ylACC1 | PYLXP' carrying GH2PS from *Gerbera hybrida*, ACC1 from *Y. lipolytica* | This study |
| PYLXP'-GH2PS-ylACC1-ylMAE | PYLXP' carrying GH2PS from *Gerbera hybrida,* ACC1, MAE from *Y. lipolytic*a | This study |
| PYLXP'-GH2PS-ylCC1-ylMAE-ylBPL | PYLXP' carrying GH2PS from *Gerbera hybrida,* ACC1, MAE, BPL from *Y. lipolytica* | This study |
| PYLXP'-GH2PS-ylMAE-Ec PDH-Ec LplA | PYLXP' carrying GH2PS from *Gerbera hybrida*, MAE from *Y. lipolytica*, Ec PDH and Ec LplA from *E coli*. | This study |
| PYLXP'-GH2PS-ylACC1-ylMAE-Ec PDH-Ec LplA | PYLXP' carrying GH2PS from *Gerbera hybrida*, ACC1, MAE from Y. lipolytica, Ec PDH and Ec LplA from *E coli*. | This study |
| PYLXP'-GH2PS-ylACC1-yltMAE | PYLXP' carrying GH2PS from *Gerbera hybrida*, ACC1, tMAE from *Y. lipolytica* | This study |
| PYLXP'-GH2PS-ylACC1 -ylMnDH1 | PYLXP' carrying GH2PS from Gerbera hybrida, ACC1, MnDH1 from *Y. lipolytica* | This study |
| PYLXP'-GH2PS-ylACC1 -ylMnDH2 | PYLXP' carrying GH2PS from *Gerbera hybrida*, ACC1, MnDH2 from *Y. lipolytica* | This study |
| PYLXP'-GH2PS-ylACC1-ylIDP2 | PYLXP' carrying GH2PS from *Gerbera hybrida*, ACC1, IDP2 from *Y. lipolytica* | This study |
| PYLXP'-GH2PS-ylACC1-ylUGA2 | PYLXP' carrying GH2PS from *Gerbera hybrida*, ACC1, ylUGA2 from *Y. lipolytica* | This study |
| PYLXP'-GH2PS-ylACC1-ylMAE-ylPDC1-ylALD4 | PYLXP' carrying GH2PS from *Gerbera hybrida*, ACC1, MAE, ylPDC1, ylALD4 from *Y. lipolytica* | This study |
| **Strains** | | |
| *E. coli NEB5α* | fhuA2 Δ(argF-lacZ)U169 phoA glnV44 Φ80 Δ(lacZ)M15 gyrA96 recA1 relA1 endA1 thi-1 hsdR17 | New England Biolabs |
| *Y. lipolytica po1g* | matA, xpr2-332, axp-2, leu2-270 | Madzak et al. |
| *HLYali101* | *Y. lipolytica po1g with vector PYLXP'-GH2PS-ylACC1-ylMAE* | This study |
| *HLYali102* | *Y. lipolytica po1g with vector PYLXP'-GH2PS-ylMAE-ylPDC1-ylALD4* | This study |

| | | |
|---|---|---|
| *HLYali103* | *Y. lipolytica po1g with vecto rPYLXP'-GH2PS-ylMAE-Ec PDH-Ec LplA* | This study |
| *HLYali104* | *Y. lipolytica po1g with vector PYLXP'-GH2PS-ylACC1-ylMAE-Ec PDH-Ec LplA* | This study |
| *HLYali105* | *Y. lipolytica po1g with integration of GH2PS-ylMAE-Ec PDH-Ec LplA into genome* | This study |
| *HLYali106* | *Y. lipolytica po1g with integration of GH2PS-ylMAE-ylACC-Ec PDH-Ec LplA into genome* | This study |

### Table S2. Primers and synthetic oligos/genes used in this study

| No. | Primer or synthetic gene | Nucleotide sequence (5' >3') |
|---|---|---|
| 1 | Gh2PS synthetic gene | atggggtcatatagcagtgatgatgtggaggtgatccgcgaggctggccgcgcacaaggactggctactatcctggccattggaaccgccacaccaccaaattgcgtggctcaagccgattacgcagactattatttccgggtaactaaatcagagcacatggtcgatctgaaggagaagtttaaaaggatttgcgaaaagactgccattaaaaagcgatatctcgccctcacggaagattatctccaggaaaacccaaccatgtgtgagttcatggctccctcattgaacgctcgccaggatctcgtggttaccggggtccccatgcttggcaaggaggccgccgtcaaagccatcgacgaatggggtttgccaaagtcaaaaattacccatctgattttctgcactaccgccggggtagacatgcccggtgcagactaccagctggtgaagctgctgggtctttccccatctgtgaagcgctatatgctgtaccagcagggctgtgcagctggtggtactgtgctgcgcctggctaaggacttggcagagaacaataaggggtcacgggtgctgatcgtctgctccgagattacagccatcctgtttcacggaccaaacgagaatcacctcgactcactggtggctcaagctctgttcggcgacggtgctgccgcgctgatagtggggtcagggccccatctggctgtggaacggcccatctttgagattgttagcacagatcagaccatcttgcccgacactgagaaagcgatgaagcttcacttgagggaggggggtctcacgttccagctccatagggacgtgccactgatggttgcaaaaaacatcgaaaacgctgccgagaaagcgctgtctccattggggattacagactggaactctgtgttttggatggttcaccctggagggagagcaatcctggatcaggtggagcgcaaactgaaccttaaagaggacaaactgagagccagcagacacgtgctgagcgagtatggaaacttgatttctgcttgcgtgctttttatcatcgacgaggtgcgcaagcgctccatggccgaaggtaagagcactaccggggaggggctggattgtggagtgctctttggatttggtccaggtatgaccgtcgaaactgttgtactccgatccgttcgagtgaccgctgcagtggccaacggcaactaa |
| 2 | ylACC1_UP synthetic gene (intron sequence is underlined) | CATCCGACCAGCACTTTTTGCAGTACTAACCGCAGCGACTGCAATTGAGGACACTAACACGTCGGTTTTTCAGTATGGCTTCAGGATCTTCAACGCCAGATGTGGCTCCCTTGGTGGACCCCAACATTCACAAAGGTCTCGCCTCTCATTTCTTTGGACTCAATTCTGTCCACACAGCCAAGCCCTCAAAAGTCAAGGAGTTTGTGGCTTCTCACGGAGGTCATACAGTTATCAACAAGGTCCTCATCGCTAACAACGGTATTGC |
| 3 | ylACC1 _F | caaggtcctcatcgctaacaacggtattgccg |
| 4 | ylACC1 _R | GGACAGGCCATGGAACTAGTCGGTACCTCACAACCCCTTGAGCAGCTCAGCCCG |
| 5 | ylMAE _F | ccgaccagcactttttgcagtactaaccgcagttacgactacgaaccatgcgaccc |
| 6 | ylMAE _R | GGACAGGCCATGGAACTAGTCGGTACCCTAGTCGTAATCCCGCACATGGATG |
| 7 | yltMAE _F | ccgaccagcactttttgcagtactaaccgcagtcctcctccagcccctccagcttcg |
| 8 | yltMAE _R | ggacaggccatggaactagtcggtaccctagtcgtaatcccgcacatggat |
| 9 | ylBPL _F | ccgaccagcactttttgcagtactaaccgcagaacgtgctggtgtataacggtccag |

| | | |
|---|---|---|
| 10 | ylBPL _R | GGACAGGCCATGGAACTAGTCGGTACCTCACTCCTTACGCTTAAGGAGTCCTC |
| 11 | ylPDH_E1aF | CCGACCAGCACTTTTTGCAGTAC**TAACCGCAG**CTCACTGCCGCTCGACGATCTAC |
| 12 | ylPDH_E1aR | GGACAGGCCATGGAACTAGTC**GGTACC**TTAGTTCTTAAAGTAGTAGTCCTC |
| 13 | ylPDH_E1bF | CCGACCAGCACTTTTTGCAGTAC**TAACCGCAG**ACTGTCAGAGACGCCCTCAACAC |
| 14 | ylPDH_E1bR | GGACAGGCCATGGAACTAGTC**GGTACC**CTACTCCTCAATGTAGAGGGCGTCC |
| 15 | ylPDH_E2F | CCGACCAGCACTTTTTGCAGTACTAACCGCAGACCCAGGGTAACATTGGCGCCTGG |
| 16 | ylPDH_E2R | GGACAGGCCATGGAACTAGTCGGTACCCTACAACAACATCTCAATGGGGTTT |
| 17 | Ec LplA _F | ccgaccagcactttttgcagtac**taaccgcag**tccacattacgcctgctcatctc |
| 18 | Ec LplA _R | GGACAGGCCATGGAACTAGTC**GGTACC**CTACCTTACAGCCCCCGCCATCCATG |
| 19 | Ec LpdA _F | ccgaccagcactttttgcagtac**taaccgcag**agtactgaaatcaaaactcaggtc |
| 20 | Ec LpdA _R | GGACAGGCCATGGAACTAGTC**GGTACC**TTACTTCTTCTTCGCTTTCGGGT |
| 21 | Ec AceF _F | ccgaccagcactttttgcagtac**taaccgcag**gctatcgaaatcaaagtaccggac |
| 22 | Ec AceF _R | GGACAGGCCATGGAACTAGTC**GGTACC**TTACATCACCAGACGGCGAATGTC |
| 23 | Ec AceE _F | ccgaccagcactttttgcagtac**taaccgcag**tcagaacgtttcccaaatgacgtg |
| 24 | Ec AceE _R | ggacaggccatggaactagtc**ggtacc**ttacgccagacgcgggttaactt |
| 25 | ylMnDH1 _F | ccgaccagcactttttgcagtactaaccgcagcctgcaccagcaacctacgctactg |
| 26 | ylMnDH1 _R | GGACAGGCCATGGAACTAGTCGGTACCTCAAGGACAACAGTAGCCGCCATC |
| 27 | ylMnDH2 _F | ccgaccagcactttttgcagtactaaccgcagtctggaccttccaccctcgccacg |
| 28 | ylMnDH2 _R | GGACAGGCCATGGAACTAGTCGGTACCTCAGAGGCAAGGTAGAGGTAGGTAG |
| 29 | ylIDP2 _F | ccgaccagcactttttgcagtac**taaccgcag**tccaccaccgctactcgaggcctg |
| 30 | ylIDP2 _R | GGACAGGCCATGGAACTAGTCGGTACCCTAAGCCAGGTCCTTCTTCAGTC |
| 21 | ylGND2_F | ccgaccagcactttttgcagtactaaccgcagactgacacttcaaacatcaagtg |
| 32 | ylGND2_R | GGACAGGCCATGGAACTAGTCGGTACCTTAAGCATCGTAAGTGGAAGAAG |
| 33 | ylUGA2 _F | ccgaccagcactttttgcagtac**taaccgcag**ttgcgagccctgaataccgtccag |
| 34 | ylUGA2 _R | GGACAGGCCATGGAACTAGTC**GGTACC**TTAAGGCTGAATGTGGGGCTCGACG |
| 35 | ylPDC1 _F | ccgaccagcactttttgcagtactaaccgcagagcgactccgaaccccaaatggtc |
| 36 | ylPDC1_R | GGACAGGCCATGGAACTAGTCGGTACCCTAAACGTTGGTCTTGGCAGAGAG |
| 37 | ylALD4 _F | ccgaccagcactttttgcagtactaaccgcagtctcttttcagcaaacttaccctag |
| 38 | ylALD4_R | GGACAGGCCATGGAACTAGTCGGTACCTTACTGCAGCTTGGCCTGGTCAAAC |

**Table S3** The retention time for the chemicals analyzed by HPLC.

| Chemicals | TAL | Glucose | Mannitol | Citric acid | Acetic acid |
|---|---|---|---|---|---|
| Retention time (min) | 23.232 | 4.977 | 5.340 | 4.595 | 7.447 |

**Table S4**. Comparison of TAL production among modified strains.

| Strains | Medium | TAL productivity (g/L) | DCW (g) | TAL to DCW yield (mg/g) | TAL to glucose/acetate yield (mg/g) | Space-time yield (mg/L h) |
|---|---|---|---|---|---|---|

| | | | | | | |
|---|---|---|---|---|---|---|
| *HLYali101* | Glucose YNB medium | 1.05 | 11.53 | 91.10 | 0.026 | 6.25 |
| *HLYali103* | | 0.81 | 8.93 | 90.71 | 0.020 | 4.82 |
| *HLYali101* | Acetate sodium YNB medium | 1.29 | 4.29 | 300.70 | 0.043 | 7.68 |
| *HLYali103* | | 1.14 | 5.52 | 206.52 | 0.038 | 6.79 |
| *HLYali101* | Glucose YNB medium in PBS buffer | 2.04 | 8.66 | 235.57 | 0.091 | 12.14 |
| *HLYali103* | | 1.96 | 7.01 | 279.60 | 0.098 | 11.67 |
| *HLYali101* | Glucose YNB medium in PBS buffer with cerulenin added | 3.20 | 7.78 | 411.31 | 0.101 | 19.05 |
| *HLYali103* | | 2.57 | 8.86 | 290.07 | 0.067 | 15.30 |
| *HLYali105* | YPD medium | 3.05 | 16.05 | 190.03 | 0.076 | 15.56 |
| *HLYali106* | YPA medium | 4.76 | 11.68 | 407.5 | 0.149 | 28.33 |

### References


Abe, I., Y. Takahashi, H. Morita and H. Noguchi (2001). "Benzalacetone synthase: a novel polyketide synthase that plays a crucial role in the biosynthesis of phenylbutanones in Rheum palmatum." European Journal Of Biochemistry **268**(11): 3354-3359.

Elmahdi, I., F. Baganz, K. Dixon, T. Harrop, D. Sugden and G. Lye (2003). "pH control in microwell fermentations of S. erythraea CA340: influence on biomass growth kinetics and erythromycin biosynthesis." Biochemical Engineering Journal **16**(3): 299-310.

Szabo, R. (1999). "Dimorphism inYarrowia lipolytica: filament formation is suppressed by nitrogen starvation and inhibition of respiration." Folia Microbiologica **44**(1): 19-24.

Szabo, R. and V. Štofaníková (2002). "Presence of organic sources of nitrogen is critical for filament formation and pH-dependent morphogenesis in Yarrowia lipolytica." Fems Microbiology Letters **206**(1): 45-50.

Xu, P., K. Qiao and G. Stephanopoulos (2017). "Engineering oxidative stress defense pathways to build a robust lipid production platform in Yarrowia lipolytica." Biotechnol Bioeng **114**(7): 1521-1530.
